# Mefloquine reduces the bacterial membrane fluidity of *Acinetobacter baumannii* and distorts the bacterial membrane when combined with polymyxin B

**DOI:** 10.1101/2025.01.15.633232

**Authors:** Nagendran Tharmalingam, Harikrishna Sekar Jayanthan, Jenna Port, Fernanda Cristina Possamai Rossatto, Eleftherios Mylonakis

## Abstract

*Acinetobacter baumannii* is a high-priority organism for the development of new antibacterial treatments. We found that the antimalarial medication mefloquine (MFQ) permeabilized the bacterial cell membrane of *A. baumannii*, decreased membrane fluidity, and caused physical injury to the membrane. MFQ also maintained activity across different pH conditions (PH range 5-8). Structure-activity relationship analysis using MFQ analogs demonstrated that piperidin-2-yl methanol is required for antibacterial activity. Scanning and transmission electron microscopy demonstrated the compromised morphological and membrane integrity in MFQ treated cells. MFQ synergized with the membrane permeabilizers polymyxin B and colistin and the MFQ+polymyxin B combination killed bacterial cells more effectively than either treatment alone. MFQ+polymyxin B was effective against other Gram-negative bacteria including *Escherisia coli, Burkholderia pseudomallei, Klebsiella pneumoniae,* and *Pseudomonas auroginosa*. Bodipy-cadaverine displacement assays confirmed the active interaction of MFQ with other membrane lipid components, such as lipopolysaccharide, lipid A, lipoteichoic acids, and fatty acids. In all-atom molecular dynamics simulations, lipid interactions facilitated the permeation of MFQ into the simulated Gram-negative membrane. Additionally, positively charged nitrogen in the piperidine group of MFQ seems to enhance interactions with the negatively charged components of the bacterial membrane. MFQ+polymyxin B caused significantly greater curvature in the simulated membrane, indicating greater damage than standalone drug treatment. Finally, *in vivo* assays showed that MFQ+polymyxin B rescued *Galleria mellonella* larvae infected with *A. baumannii*. In conclusion, membrane-active agents such as MFQ may warrant further investigation as potential component of Gram-negative infection treatment, particularly in combination with polymyxin B.

**Importance:** Antimicrobial resistance is a threat globally, and new treatments are urgently needed to combat the rise of multidrug-resistant bacteria. However, the development of anti-infectives has declined over the last two decades due to regulatory, financial and long-term requirement related challenges. In this study, we examined the membrane interactions of the antiparasitic agent mefloquine in combination with polymyxin B, using both *in vitro* and *in silico* approaches to evaluate their potential efficacy against Gram-negative bacterial infections. We investigated the interaction of MFQ with lipid bilayers to understand the mechanism through which antibacterial activity is exerted. The piperidine moiety of MFQ plays a critical role in its interaction with the lipid bilayer and facilitates membrane permeabilization. In contrast, the membrane permeabilizer polymyxin B is associated with significant neurotoxicity and nephrotoxicity. Our findings highlight the potential of membrane-acting compounds, such as MFQ, to enhance combinatorial activity while mitigating polymyxin B-associated toxicity.

## Introduction

*Acinetobacter baumannii* is a nonflagellated, coccobacillus-shaped, aerobic Gram-negative bacterium characterized by significant genomic and phenotypic diversity among its isolates (1). The adaptable genetic machinery enables variable virulence as well as the rapid development of resistance and survival mechanisms tailored to the environment (2, 3). *A. baumannii* is part of the ESKAPE group (4), which includes six critical multidrug-resistant pathogens and is linked to severe hospital-acquired infections, exploiting wounds and invasive procedures (like catheters and ventilation) to life-threatening conditions (5). The World Health Organization has listed *A. baumannii* as a critical-priority pathogen for the development of new treatments (6), while the Centers for Disease Control and Prevention (CDC) has classified *A. baumannii* as an urgent public health threat (7).

Virulence factors of *A. baumannii* include lipopolysaccharides, capsular polysaccharides, proteases, outer membrane porins, biofilm-associated proteins, iron-chelating systems, glycoconjugates, and protein secretion systems (3). Also, *A. baumannii* exhibits antibiotic resistance (8) by overexpressing efflux pumps (e.g., AdeABC), producing β-lactamases (e.g., carbapenemases) (9), mutating or losing porins (e.g., CarO) (10), and using regulatory systems (e.g., AdeRS) to enhance resistance gene expression (2). Target site modifications, such as mutations in DNA gyrase or ribosomal targets (11), hinder drug binding, while enzymes like aminoglycoside-modifying enzymes (AMEs) chemically alter antibiotics (12).

*A. baumannii* bacteria also form biofilms on surfaces, which not only create physical barriers to drugs, but also facilitate the acquisition of resistance genes through plasmids, transposons, or integrons (13). Biofilm structures act as a protective barrier that enhance resistance by limiting antibiotic penetration, evading the immune system and enabling persistence in harsh environments (14). The protective nature of biofilms promotes chronic infections, and increase the expression of resistance-associated genes further complicating treatment (15). Consequently, biofilm-related infections, such as ventilator-associated pneumonia and catheter-associated infections, pose significant clinical challenges (16), often requiring combination therapies and higher antibiotic doses.

Through a large screening, we discovered that the anti-malarial mefloquine (MFQ) is active against *Francisella tularensis* (17). *F. tularensis* is a Gram-negative pathogen, hence we hypothesized that MFQ can inhibit other Gram-negative pathogens using broth-microdilutions assays and found that the MFQ is active against *A. baumannii*. While we were pursuing our investigation, the antibacterial activity of MFQ against Gram-negative pathogens was reported (18, 19). Here, we confirm and expand upon these observations by studying the molecular membrane interaction behavior of MFQ in combination with polymyxin B, as polymyxin B is a potent antibiotic effective against Gram-negative bacterial infections. We report the antibacterial properties of MFQ against *A. baumannii* through membrane-permeabilizing action and provide a detailed understanding of the membrane interactions in the MFQ-polymyxin B combination.

## Results

### Antibacterial susceptibility

Evaluation of MFQ against a panel of ESKAPE pathogens using broth microdilution assays showed a minimum inhibitory concentration (MIC) of 32 mg/L against *A. baumannii* (**Fig. 1A**). MFQ inhibited other *Acinetobacter* spp. (**Table 1**) and killing kinetics assays showed that MFQ at 2x and 4x MIC caused a 4 log_10_-fold decrease in colony-forming units (CFU)/mL from the initial inoculum compared to DMSO controls (**Fig. 1B**), indicating bactericidal activity.

**Figure 1.**
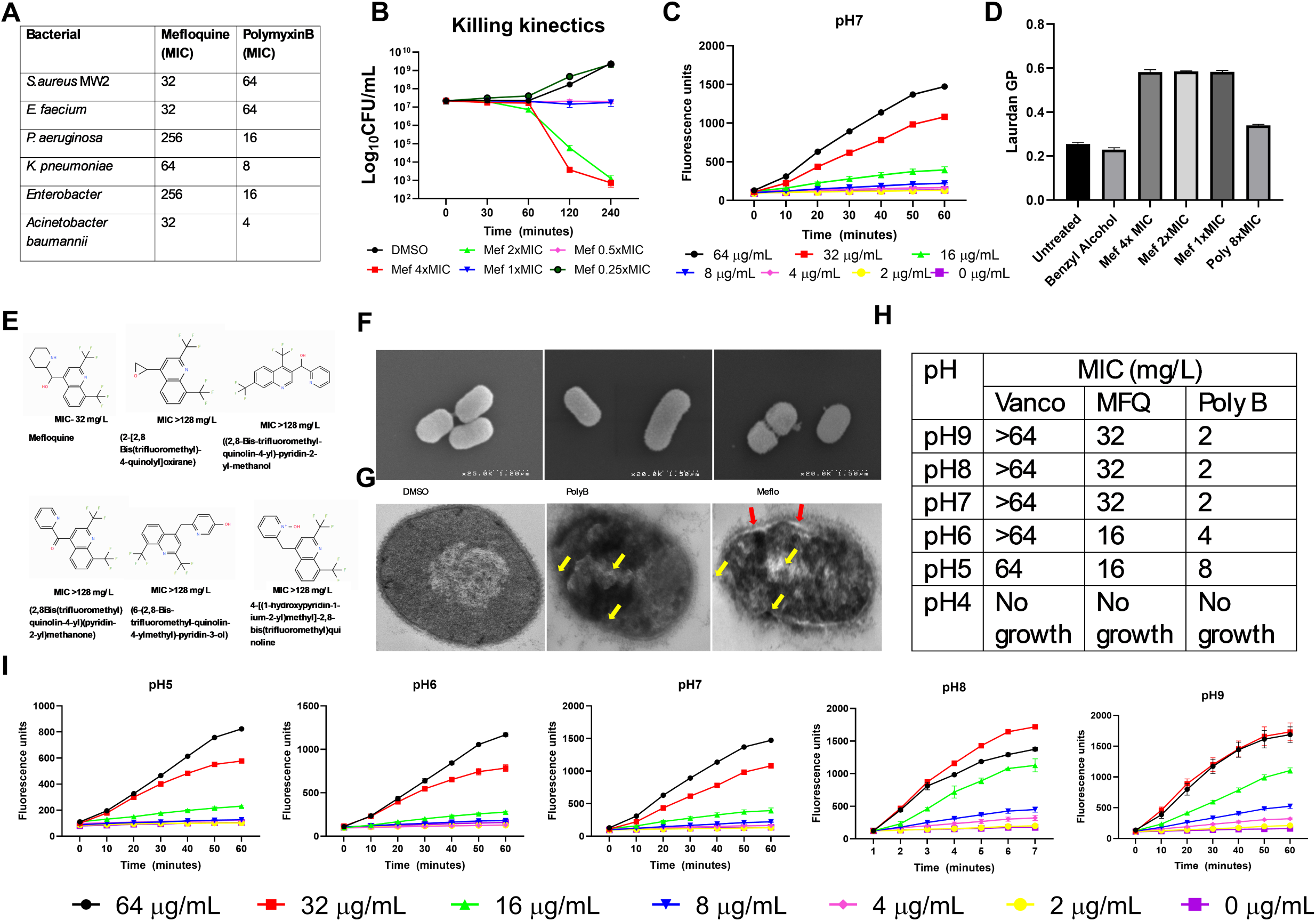
Antibacterial activity of mefloquine against *A. baumannii*. **A.** A broth microdilution assay was used to determine the antibacterial susceptibility of major nosocomial ESKAPE pathogens to MFQ and polymyxin B. **B**. Killing kinetics assays were used to evaluate the bactericidal activity of mefloquine (MFQ). Exponentially growing *A. baumannii* 17978 at 5x10^7^ cells were treated with various concentrations of MFQ or dimethyl sulfoxide (DMSO) (negative control), and colony-forming units (CFU)/mL were counted at various intervals. **C.** Exponentially growing *A. baumannii* 17978 were treated with various concentrations of MFQ, and cellular fluorescence was measured spectroscopically by monitoring the uptake of the live cell membrane-impermeable fluorescent dye Sytox Green (excitation = 435 nm, emission = 530 nm) during a 1-hour incubation period. **D.** Exponentially growing *A. baumannii* 17978 cells were allowed to reach late-exponential phase, Laurdan fluorescence probe-tagged cells were treated with various concentrations of MFQfor 1 hour, and the Laurdan generalized polarization (GP) was calculated as (I_440_-I_490_)/(I_440_+I_490_) based on cellular fluorescence when excited at 350 nm. *A-D* shows data with means ± SD for n = 3 biologically independent samples. **E.** Structure-activity relationship analysis of commercially available mefloquine analogs using the broth microdilution method. Compared to MFQ, compound 2 has an oxiranyl group replacement of 2-piperidinyl group, compound 3 has a 2-pyridinyl group replacement of 2-piperidinyl group, compound 4 has a 2-pyridinylcarbonyl replacement of hydroxylmethyl-(2-piperidinyl) group, compound 5 has a hydroxypyridinyl group replacement of 2-piperidinyl group and compound 6 has a N-oxide-pyridinyl replacement of 2-piperidinyl group (which is a dipolar form of pyridine). This structure includes a hydroxyl group and is more basic than a neutral pyridine. MICs are shown as the means of n = 3 biologically independent samples. **F-G.** Electron microscopy demonstrates malformation/membrane damage and blisters in the MFQ-treated cells. *A. baumannii* 17978 cells were treated MFQ at 4x MIC for 30 minutes, washed, fixed, and analyzed using scanning electron microscopy (**F**), and resin-imbedded cells were visualized with transmission electron microscopy (**G**). **H.** Broth microdilution assay was used to determine the antibacterial susceptibility of *A. baumannii* 17978 under different pH conditions. MICs are shown as the means of n = 3 biologically independent samples. **I.** Exponentially growing *A. baumannii* 17978 were treated with various concentrations of MFQ, the pH of the medium was adjusted with sterile NaOH or 1N HCl, and membrane permeability was measured as described in C. All graphs show data with means ± SD for n = 3 biologically independent samples.

**Table 1.** Antibacterial activity of MFQ against various *Acinetobacter* spp. A broth microdilution assay was used to determine the antibacterial susceptibility of *Acinetobacter spp*. pathogens to MFQ and polymyxin B. MICs are shown as the means of n = 3 biologically independent samples.

| Bacteria | Minimum inhibitory concentration (MIC; mg/L) |  |
| --- | --- | --- |
|  | Mefloquine | PolymyxinB |
| *A. anitratus* | 32 | 8 |
| *A. iwoffii* | 8 | 0.5 |
| *A. hemolytica* | 16 | 8 |
| *A. boydii* | 16 | 4 |
| *Acinetobacter calcoaceticus* | 32 | 8 |
| *A. baumannii* HKU 4751 | 32 | 2 |
| *A. baumannii* 307 | 64 | 8 |
| *A. baumannii* 0059 | 64 | 8 |
| *A. baumannii* 19606 | 32 | 4 |
| *A. baumannii* 900 | 32 | 8 |
| *A. baumannii* M2 | 32 | 16 |

Next, we conducted a series of studies to investigate MFQ-membrane interactions *in vitro*. We first performed a membrane permeability assay using a membrane-impermeable, DNA-binding fluorescent dye Sytox Green. The fluorescence assay showed an increase in cellular fluorescence in cells treated with MFQ at 4x or 1x MIC compared to DMSO (**Fig. 1C**) indicating that MFQ permeabilizes the bacterial cell membrane. Then, we tested membrane fluidity using the Laurdan membrane-embedding fluorescent dye. The generalized polarization (GP) values, calculated from cellular fluorescence, revealed higher GP values in cells treated with MFQ at 1x, 2x, and 4x MIC compared to untreated cells and benzyl alcohol-treated controls (**Fig. 1D**), indicating that MFQ decreases membrane fluidity.

To validate the critical component of MFQ as an antibacterial agent, we studied structure-activity relationship analysis. We used commercially available MFQ analogs that are lacking piperidine ring connected to the 4-position of the quinoline ring system through a secondary alcohol group to test antibacterial activity. We tested MFQ analogs using a broth microdilution method showed a complete loss of antibacterial activity (MICs > 128 mg/L) **(Fig. 1E**) compared to the parental MFQ molecule.

We used scanning electron microscopy (SEM) (**Fig. 1F**) and transmission electron microscopy (TEM) to visualize the structural integrity of *A. baumannii* treated with MFQ, (**Fig. 1G**). We found significant morphological abnormalities include altered cell-shape and cellular structure, uneven surface textures in SEM and MFQ-induced membrane damage in TEM. Notable observations included disrupted membrane integrity, highlighted by the presence of an expanded inner membrane space (**Fig. 1G**; *red arrow*), and a loss of structural uniformity (**Fig. 1G**; *yellow arrow*).

We also tested the antibacterial potential of MFQ at various pH levels and found that MIC of MFQ dropped from 32 mg/L to 16 mg/L at pH 6 and 5 (**Fig. 1H**). In membrane permeability assays under various pH conditions, MFQ treatment showed an increase in cellular Sytox Green fluorescence, indicating that MFQ remained active as a membrane permeabilizer at acidic pH (**Fig. 1I**). A mutation frequency assay revealed that MFQ inhibited *A. baumannii* over a 100-day period of continuous culture, whereas rifampicin yielded resistance development (**Supplementary Figure 1**).

Next, we evaluated the anti-biofilm activity of MFQ, focusing on the ability to inhibit biofilm formation and eradicate matured biofilms. MFQ treatment of *A. baumannii* resulted in a significant reduction of at least 50% in the optical density of biofilms at concentrations ranging from 16 to 128 mg/L (**Fig. 2A**; 4-0.5 × MIC). We also evaluated the efficiency of MFQ in eradicating matured biofilms and observed a reduction of at least 50% in the optical density of matured biofilms at concentrations ranging from 64 to 128 mg/L (**Fig. 2B**; 4×-2× MIC). Subsequent real-time polymerase chain reaction analysis revealed that genes associated with biofilm, such as the *csuA, csuB, csuC* and *pgaA, pgaB*, *pgaC* were downregulated after treatment with sub-MIC levels of MFQ (**Fig. 2C**). In addition, lipopolysaccharide-associated genes, such as *pmrABC*, were also downregulated (**Fig. 2C**), indicating that MFQ treatment is associated with downregulation of genes involved in pili and lipopolysaccharide formation.

**Figure 2.**
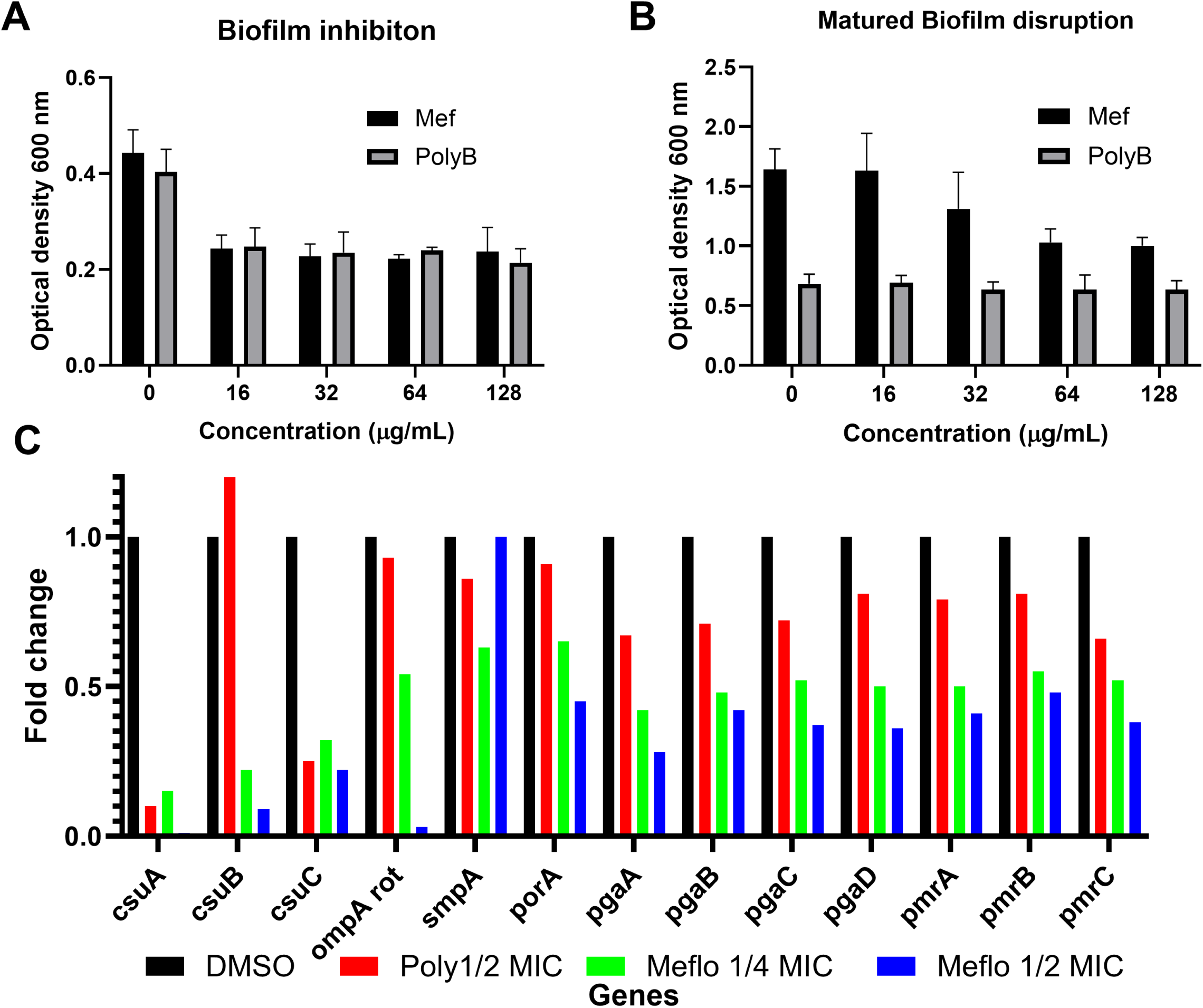
Biofilm studies. MFQ inhibit biofilm formation (***A***) and disrupts mature biofilm (***B***). To monitor biofilm formation, glucose-supplemented *A. baumannii* 17978 cells were grown in the presence of MFQ at various concentrations. After 24 h of incubation, the wells were washed three times, MHB supplemented with the Alamar Blue HS reagent was added, and the metabolic activity was measured in a spectrometer at 570 nm. All graphs show data with means ± SD for n = 3 biologically independent samples**. C.** MFQ down-regulates biofilm-assocaited genes. *A. baumannii* 17978 were treated with sub-MIC of MFQ and total RNA was isolated. The biofilm-associated genes were probed, and gene regulation was monitored using real-time PCR. Fold change was calculated from triplicate values.

### MFQ synergized with the membrane permeabilizer antibacterial agent polymyxin B

We performed a series of checkerboard assays and found that the MIC of MFQ decreased from 32 to 8 mg/L in the presence of polymyxin B at 2 mg/L (**Fig. 3**) with the fractional inhibitory concentration value of 0.5 indicating synergism. Next, we extended testing of MFQ+polymyxin B combination against other Gram-negative pathogens, such as *Escherichia coli, Burkholderia pseudomallei, Klebsiella pneumoniae, Pseudomonas aeruginosa,* and *Enterobacter aerogenes.* Checkerboard assays demonstrated a 2-log reduction in the MIC of MFQ when used in combination, compared to standalone drug treatment. (**Fig. 3**). We also included colistin in the checkerboard assays and found a 2-log reduction in the MIC of MFQ when used in combination, compared to standalone drug treatment (**Fig. 3**).

**Figure 3.**
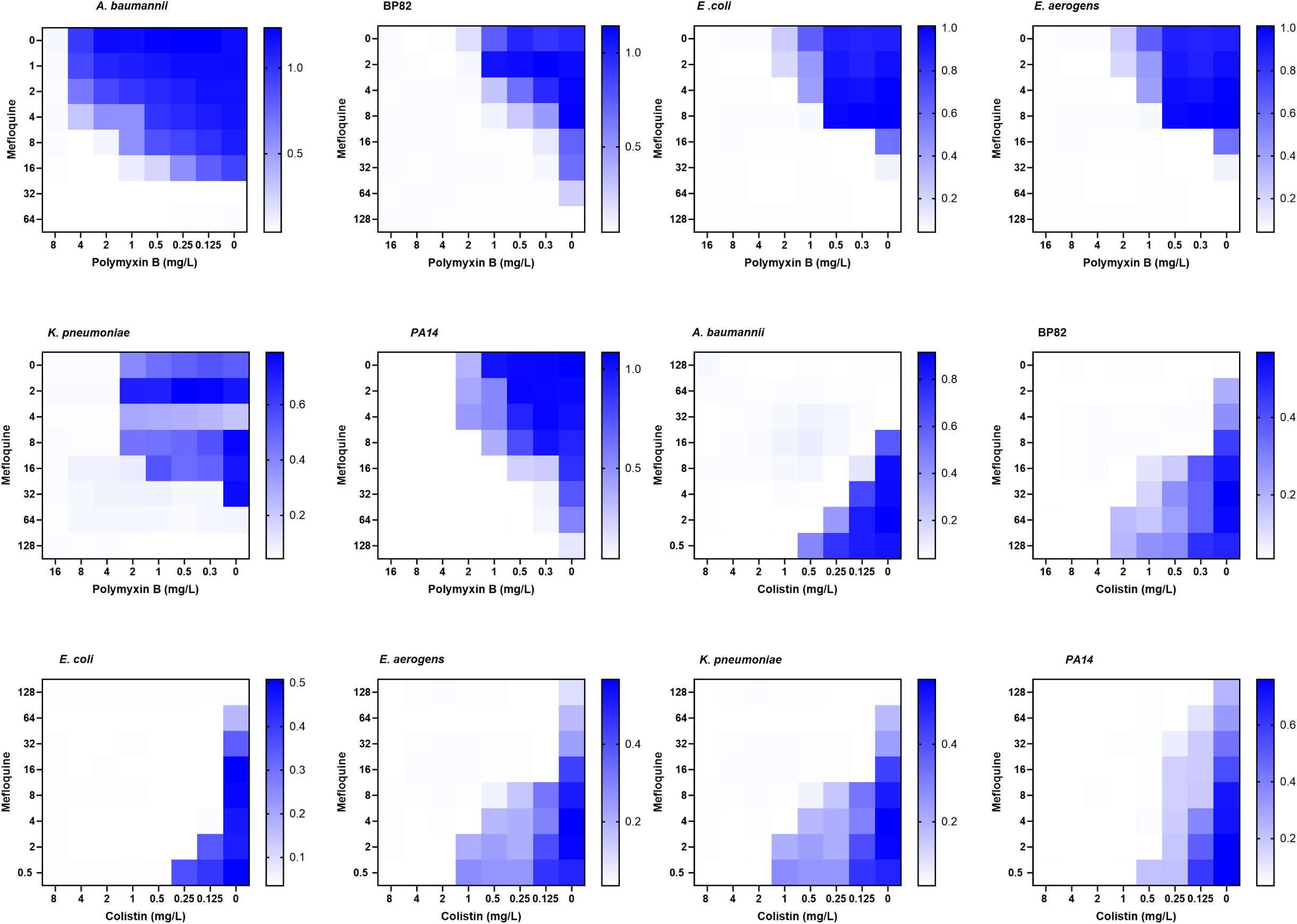
MFQ is active against *A. baumannii* and other Gram-negative bacteria in the presence of Polymyxin B or colistin. Combined treatment of MFQ and a lipopeptide polymyxin B or colistin was evaluated using broth microdilution checkerboard assays against *A. baumannii, K. pnaemoniae, P. aurenoginosa, E. aerogens E. coli*, *B. pseudomallei* (BP82; BSL2). The fractional inhibitory concentration index (FICI) was determined from the optical densities of drug combinations evaluated at 600 nm after an 18-h incubation at 37°C (synergy: FICI ≤ 0.5; no interaction: 0.5 < FICI ≤ 4; antagonism: FICI > 4). Individual biologically independent experiments were carried out in triplicate and representative results are shown.

We also evaluated the potential of MFQ+polymyxin B combination using killing kinetics, where the combination of 0.5× MIC of MFQ and 0.5× MIC of polymyxin B killed 3-log_10_ CFU/mL of *A. baumannii* from the initial inoculum (**Fig. 4A**). However, standalone drug treatment of MFQ (0.5× MIC) did not reduce bacterial load, whereas polymyxin B (0.5× MIC) decreased only a 1-log_10_ CFU/mL (**Fig. 4A**). Polymyxin B at 4× MIC was used as a positive control. We further tested the effect of MFQ+ polymyxin B combination at 10× MIC concentration using TEM. The MFQ+ polymyxin B combination caused significant membrane damage resulting in severe structural perturbation compared to the DMSO control (**Fig. 4B**). The combinatorial activity of MFQ with polymyxin B prompted us to evaluate the synergy with another lipopeptide antibiotic, colistin, against *A. baumannii*. Our findings revealed that the MIC of MFQ decreased from 32 mg/L to 8 mg/L in the presence of colistin at a concentration of 2 mg/L (**Fig. 4C**)

**Figure 4.**
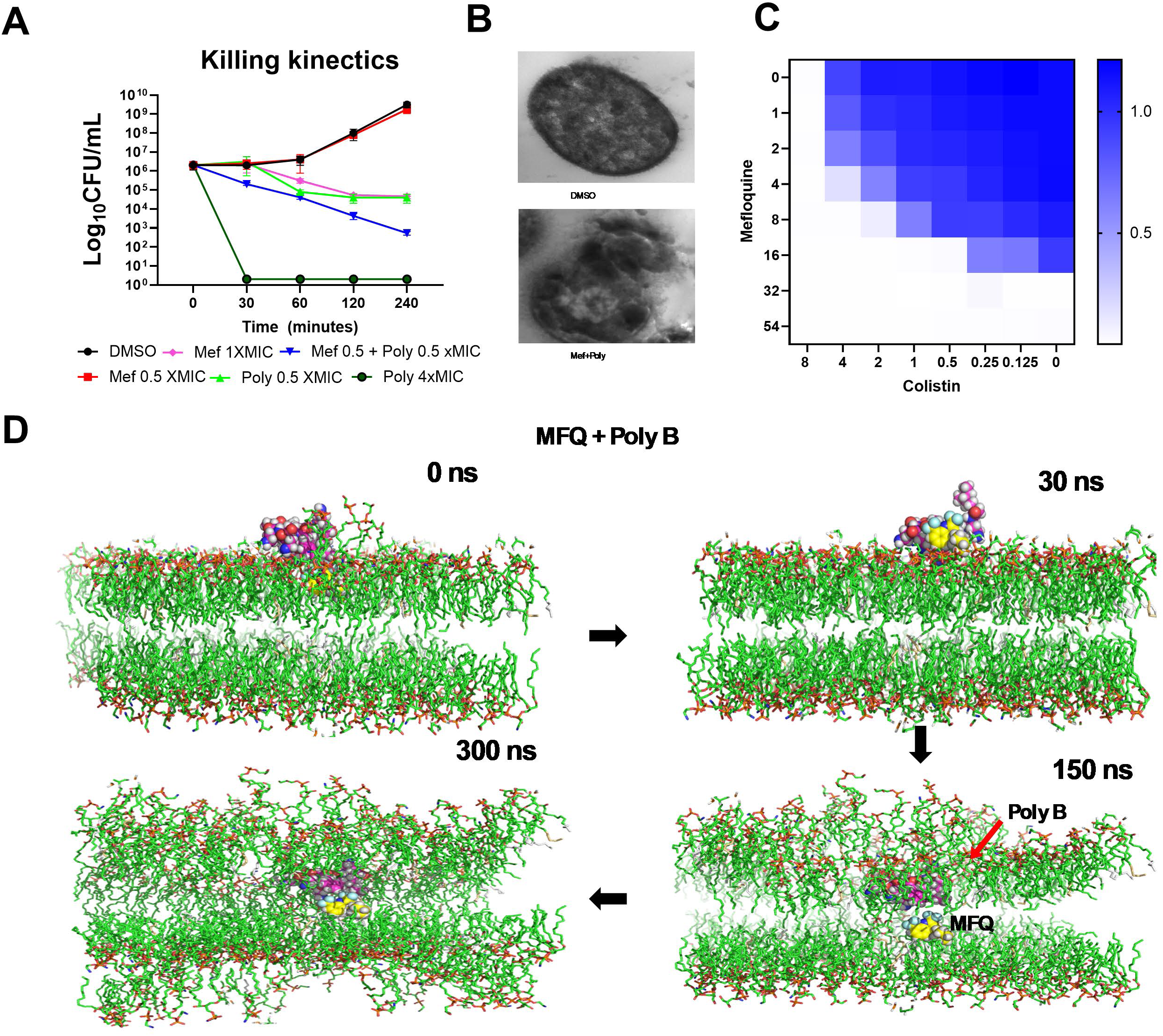
MFQ synergizes with lipopeptide antibiotics against *A. baumannii*. ***A.*** MFQ is a bactericidal agent when used in combination with polymyxin B against exponential-phase *A. baumannii* 17978 cells. Total viable counts were quantified using the spot-plate method. The graphs show data with means ± standard deviation (SD) for n = 3 biologically independent samples. ***B.*** Transmission Electron Microscopy shows malformation/membrane damage in the MFQ treated cells. *A. baumannii* 17978 cells were treated MFQ at 4xMIC for 30 minutes; fixed and resin imbedded cells were visualized under transmission electron microscopy. ***C.*** MFQ synergizes the lipopeptide antibiotic colistin. Synergistic activity between MFQ and colistin against *A. baumannii* was evaluated using broth microdilution checkerboard assays. The fractional inhibitory concentration index (FICI) was determined from the optical densities of drug combinations evaluated at 600 nm after an 18-h incubation at 37°C (synergy: FICI ≤ 0.5; no interaction: 0.5 < FICI ≤ 4; antagonism: FICI > 4). Individual biologically independent experiments were carried out in triplicate and representative results are shown. **D.** MFQ combined with polymyxin B damages simulated *A. baumannii* membranes. MD simulation studies for ∼ 1 μs of MFQ+polymyxin B interacting with a simulated *A. bauumannii* bacterial membrane bilayer were carried out at pH7. The four panels, left to right, show sequential attachment, embedding, penetration, and membrane damage, respectively. The simulation was repeated three times with similar results, and frames from a representative simulation are shown.

### Molecular dynamics (MD) simulations investigating the action of MFQ and polymyxin B on the bacterial membrane

We used all-atom ∼ 1 μs-MD simulations with lipid14 amber force field (20) to investigate the combinatorial action of MFQ and polymyxin B on a simulated *A. baumannii* like-membrane consisting of phosphatidylethanolamine (PE), phosphatidylglycerol (PG), acyl-phosphatidylglycerol (aPG) and cardiolipin (CL) together with monolysocardiolipin (MLCL) at pH 7.

Initial analysis focused on lipid packing defects by calculating the size distribution probability (membrane thickness, nm) before and after polymyxin B insertion. The analysis showed no significant differences between the pure membrane and the polymyxin B-bound membrane, suggesting that polymyxin B binding does not induce lipid packing defects. Additionally, polymyxin B binding did not cause membrane bending, alter lipid packing, or affect water permeability (data not shown). Similarly, the combination of MFQ and polymyxin B effectively helps MFQ to permeate membrane In our simulations polymyxin B, when combined with MFQ, did not permeate the membrane. (**Fig. 4D**). Further simulations showed that the piperidine ring of MFQ initially dipped into the PE/PG membrane, while the naphthyridine ring remained oriented toward the water layer, marking the initial surface binding of the molecule (**Fig. 5A).** While the drug exists predominantly in a protonated form at pH 7, it may undergo deprotonation upon crossing the membrane. For modeling purposes, MFQ was assumed to be protonated at pH 7; however, its charge was kept neutral to simulate the full permeation event. This approach accounts for the energetic cost that charged small molecules typically incur when becoming neutral to permeate the membrane via free diffusion. We also considered a potential shift in the pKa of the MFQ molecule, resulting in neutralization within the hydrophobic core region of the membrane (21). Partial charges were determined using Gaussian quantum mechanical (QM) calculations, and electrostatic potential (ESP) charges were fitted using multi-conformational charge analysis. To facilitate the simulation, the charge of MFQ was manually set to zero. Additionally, it is important to note that during the binding event with the membrane head group, charged molecules tend to form stronger electrostatic interactions, which may influence their initial interactions with the membrane. Following the initial interaction, the MFQ molecule stabilized approximately 9 to 10 Å above the center of the lipid bilayer (z = 0 Å) and rapidly inserted into the lipid acyl chains of the monolysocardiolipin/cardiolipin (MLCL/CL) membrane. Notably, the hydrophobic CF₃ group on the naphthyridine ring was the first to interact with the inner lipid surface, primarily through hydrophobic interactions, facilitating deeper integration into the bilayer.

**Figure 5.**
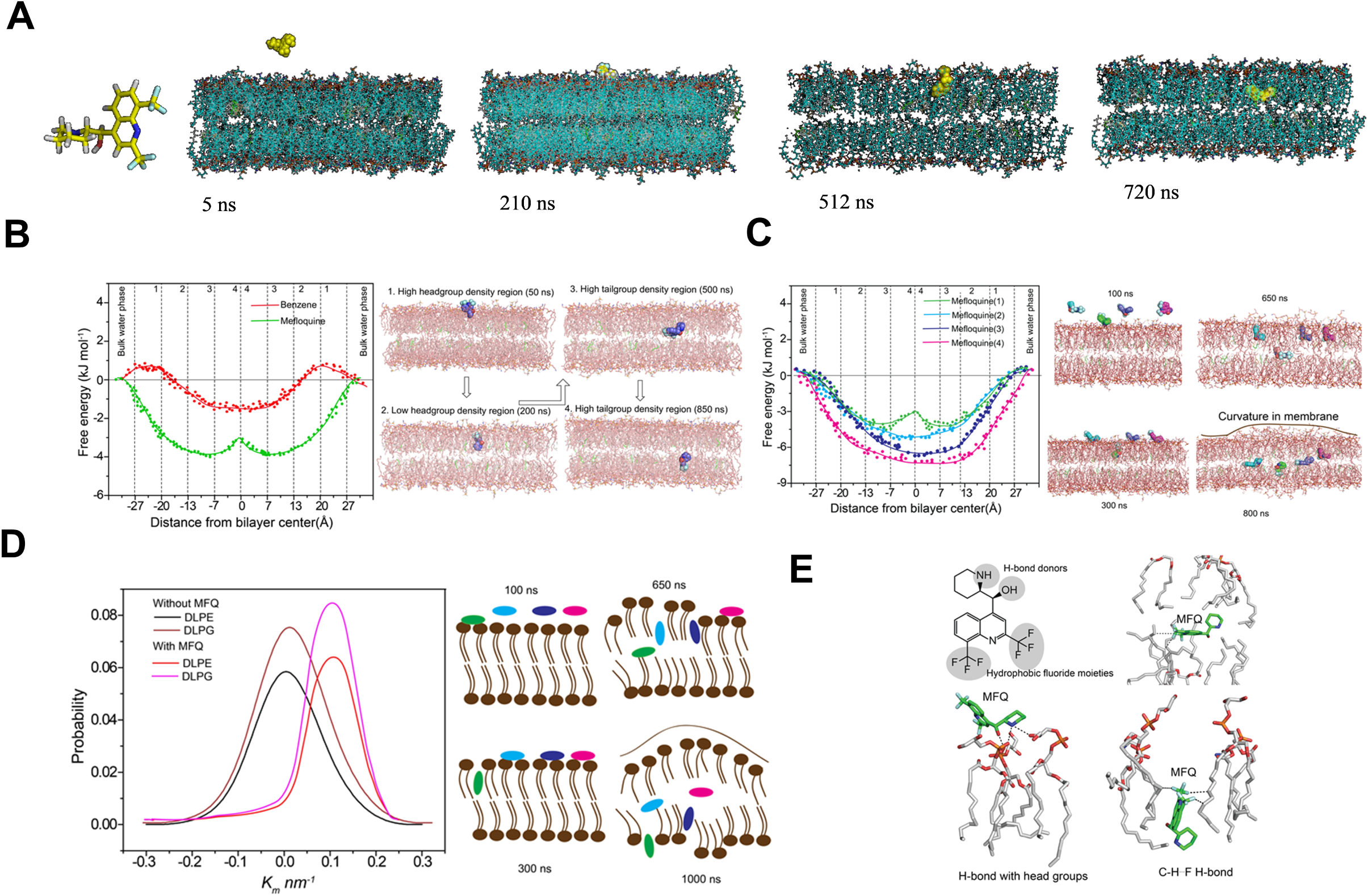
All atom molecular dynamics simulation. **A.** *In silico* analysis of membrane permeability of MFQ against Gram negative membrane at pH 7. **B**. Free energy profiles for MFQ in multi-drug-membrane system (4:1). Profiles were estimated completely across the bilayer. Each point indicates free energy, multiple points were extracted from three different simulations. MD snapshots depict the location of the drugs in the membrane during MD simulations. **C**. Free energy profiles for MFQ in multi-drug-membrane system, i.e. bilayer containing 1-4 MFQ molecules. Profiles were estimated completely across the bilayer. Each point indicates free energy, multiple points were extracted from three different simulations. **D**. Distribution of mean curvatures for lipid domains from the last 300 ns of the MD simulations shows that the mean curvature in the presence of MFQ increased to approximately 0.1 nm⁻¹. Cartoon representation of the drug induced membrane curvature. **E**. Non-covalent interactions between drug and lipid bilayer observed during the course of MD simulations. Hydrogen-bonding interactions between CH^…^F hydrogen bonds were validated using DFT calculations.

Once entered the low tail group density region, MFQ was reoriented into a horizontal plane, likely due to the formation of strong hydrophobic interactions between MFQ and the tail groups. However, the single MFQ molecule and membrane system did not exhibit any significant membrane perturbation. However, in independent simulations, multiple MFQ molecules were pre-adsorbed onto the surface of the membrane, where they altered their orientations during the simulation before positioning just before the polar head groups of the lipid bilayer. One MFQ molecule subsequently penetrated the hydrophobic region of the PE/PG membrane, stabilizing at 9.2 Å above the center of the lipid bilayer. (**Fig 5B**).

To further investigate the impact of drug-induced membrane perturbations, three additional MFQ molecules were randomly positioned on the membrane after removal of water molecules and MD simulation was resumed following water molecule equilibration (**Fig 5C**). The PMF free energy results from the multiple drug-membrane system showed that the permeation of single MFQ molecule facilitates the permeation of additional MFQ molecules. Structural perturbations in the membrane, including membrane curvature were observed. **Figure 5C** illustrates the resulting asymmetric behavior within the membrane bilayer. The distribution of mean curvature of the membrane without any drug molecules was approximately 0 nm⁻¹, while the mean curvature with the drug became approximately 0.1 nm⁻¹ due to structural perturbations of membrane in the presence of MFQ (**Fig. 5D**).

### MFQ-lipid interactions- *in silico* analysis

To investigate the interaction between MFQ and the lipid surface, we calculated the number of hydrogen bonds (H-bonds) formed between MFQ and the lipid head groups. Triplicate MD simulations of MFQ revealed that MFQ primarily interacts with the polar head groups of the PE/PG/CL membrane through H-bond formation. On average, MFQ forms at least two H-bonds during the simulations (**Table 2**). Notably, as the number of MFQ molecules increases, the number of H-bond interactions between MFQ and the lipid also increased. In the presence of MFQ, the number of H-bonds between water and the lipid head groups was approximately 775 compared to the absence of MFQ. The membrane becomes more disordered and exposed to the solvent, which facilitates additional hydrogen bond formation with water molecules (**Fig. 5E**). Free energy analyses from the simulations indicated that MFQ permeation into the lipid bilayer is highly favorable, with the minimum free energy observed at the center or tail regions of the bilayer. During the simulations of multiple MFQ molecules, once a single MFQ molecule crosses the bilayer membrane, the membrane adjusts to allow more molecules to permeate. We did not observe any further reduction in the number of molecules. Initially, MFQ molecules were positioned in a vertical orientation during free energy analyses, with the carbonyl and NH groups directed toward the membrane center, forming multiple H-bonds with the lipid head groups (**Fig. 5E**). Meanwhile, the fluoride group was oriented toward the aqueous phase and H-bond interactions were consistently observed across multiple simulations, even when starting from random orientations and positions of MFQ within the system (**Fig. 5E**). DFT calculations were used to analyze the MM-derived interactions between MFQ and the individual membrane system at various time points. The positively charged MFQ piperidine forms electrostatic interactions with the phosphatidyl head region, while the hydroxyl group in MFQ participate along with the above-mentioned electrostatic interactions as observed in the MD simulations. At the permeation stage we observe that the fluoride hydrophobic moieties form hydrophobic interactions with the lipid hydrophobic tail.

**Table 2.** Average number of different hydrogen bonds in the simulated systems: Hydrogen bonding interactions observed in ∼ 1μs-MD simulations with systems containing 1-4 MFQ molecules per membrane bilayer (BL), detailing the number and types of hydrogen bonds formed between the molecules and the surrounding environment; a system with benzene/BL was used as reference to account for the differences in H-bonds between the BL and water molecules.

| Model | No of drugs | No of H-bonds between<br>drug and water | No of H-bonds between<br>membrane and water |
| --- | --- | --- | --- |
| Benzene/BL | 1 | 0 | 778.2 ± 2.8 |
| MFQ/BL | 1 | 1.48 ± 0.1 | 782.1 ± 2.8 |
| 2MFQ/BL | 2 | 2.7 ± 0.2 | 814.6 ± 4.1 |
| 3MFQ/BL | 3 | 4.5 ± 0.4 | 837.4 ± 3.9 |
| 4MFQ/BL | 4 | 5.2 ± 0.4 | 844.8 ± 3.4 |

### MFQ -Lipid interactions- *in vitro* analysis

The aromatic rings in MFQ, including the quinoline and other aromatic components, enhance the lipophilic nature of MFQ (**Fig. 5E**). To further investigate the interaction of MFQ with membrane components, we used the fluorescent probe Bodipy cadaverine (BODIPY-Boron-Dipyrromethene cadaverine) to study binding to lipopolysaccharides (LPS). In the presence of MFQ, the displacement of cadaverine from LPS resulted in increased fluorescence intensity, indicating the extent of MFQ-LPS interaction (**Fig. 6A**). Polymyxin B, used as a positive control, produced a similar fluorescence increase, while the absence of any drug had no effect. We also examined the interaction of MFQ with Lipid A, another major membrane component of *A. baumannii*. We found that MFQ binds to Lipid A (**Fig. 6B**), and checkerboard assays further quantified MFQ+Lipid A interaction, demonstrating an increase in fluorescence units that reflects MFQ’s strong affinity for Lipid A (**Fig. 6C**).

**Figure 6.**
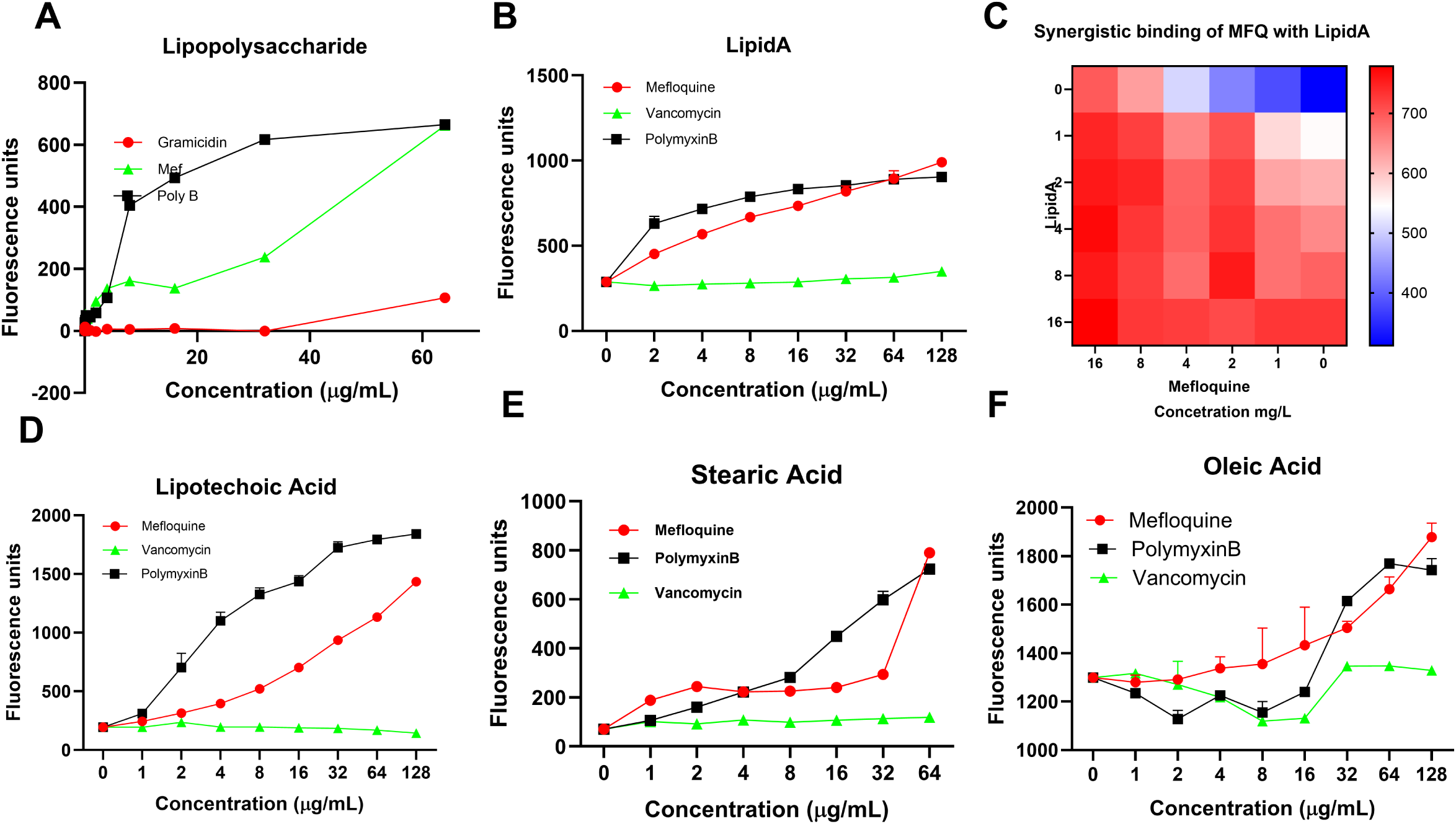
MFQ is lipophilic. The lipophilic nature of MFQ was tested using a Bodipy-cadaverine displacement assay. Bilayer components (***A***-lipopolysaccharides; ***B***-LipidA, ***D***-lipoteichoic acids; ***E***-Strearic acid, F-Oleic acid and ***C***-combination of MFQ and LipidA were added to the probe and the complex was added to serially diluted CC in Tris-HCl buffer. Fluorescence was measured at ex 590, em 620. All graphs show data with calculated means ± SD for n = 3 biologically independent samples.

### *In vivo* efficacy

First, we tested the toxicity of MFQ and found that MFQ did not show hemolysis against human red blood cells (RBCs). As a control we used polymyxin B that did not show hemolysis up to 64 mg/L, while Triton X and gramicidin A, were used as positive controls, resulting in substantial lysis (Upper X-axis-concentration of MFQ and Gramicidin. Lower X-Axis-percentage of Triton-X) (**Fig. 7A**).

**Figure 7.**
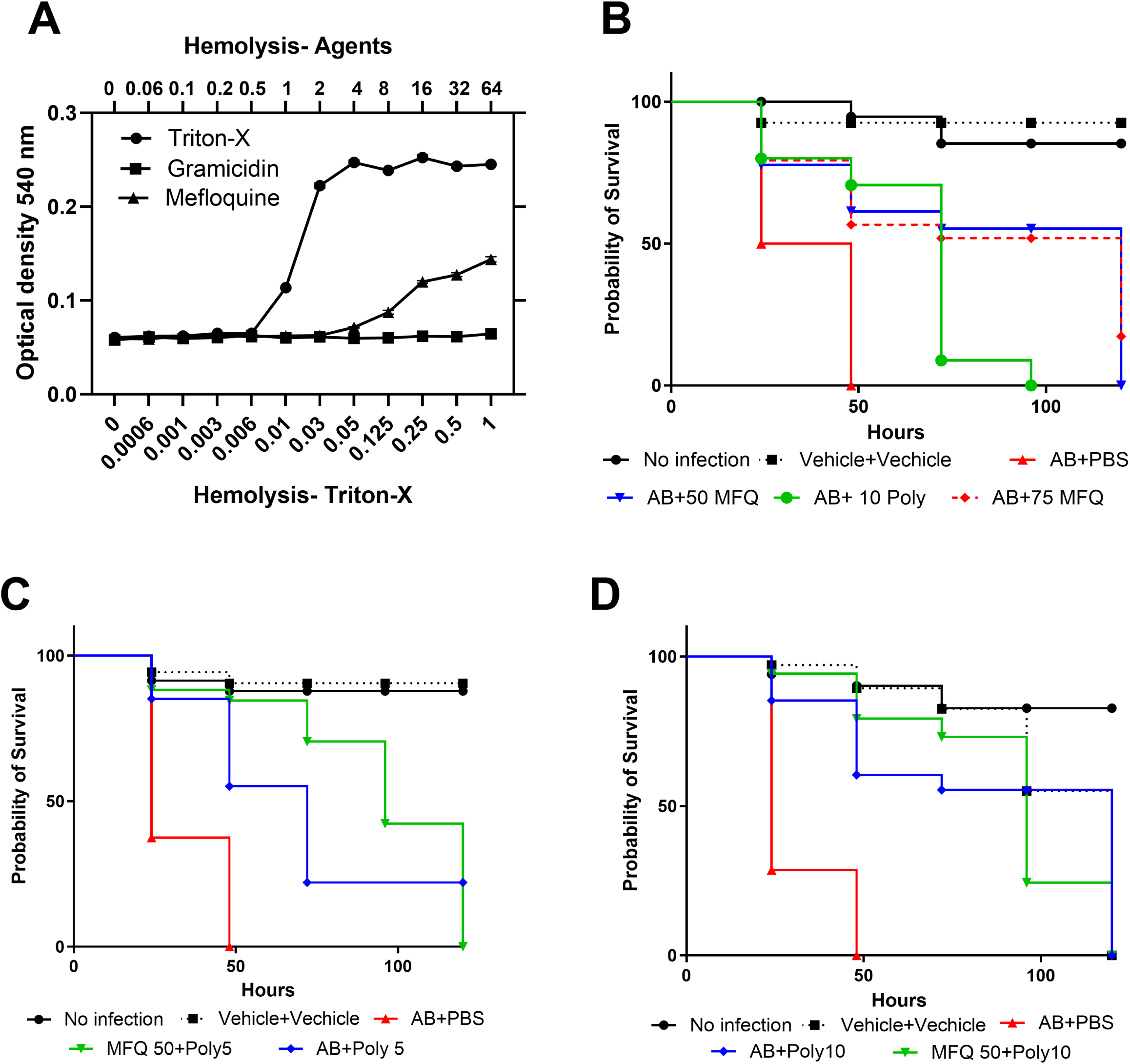
Cytotoxicity and *in vivo* efficacy of MFQ. **A.** Hemolysis potential of MEF. Human RBCs were suspended in the presence of MFQ or gramicidin or Triton-X for 60 minutes. The optical density of supernatant was measured spectroscopically at 540 nm. Data show means ± SD for n = 3 biologically independent samples. **B**. MFQ is an efficacious in an insect model. *Galleria mellonella* larvae (n = 16 biologically independent animals) were injected with *A. baumannii* 17978 in the last left proleg. After 30 minutes of infection, MFQ (5% Kolliphor + 5% Ethanol in USP Saline) or polymyxin B or combination (**C-D**) was injected into the last right proleg using a Hamilton syringe. Experiment **C&D** used the same control group. The assay was repeated three times, and representative data are shown (*p*<0.0001). Statistical significance was assessed using the Mantel-Cox log-rank test in GraphPad prism.

Next, we tested the *in vivo* efficacy of MFQ or in combination with polymyxin B using the insect *Galleria mellonella* wax moth larvae model. The larvae were injected with a lethal dose of *A. baumannii* 17978, followed by an injection of MFQ, polymyxin B, or their combination. Untreated larvae showed 100% mortality within 48h of infection. The LT50 of untreated larvae was less than 24h. However, MFQ at 50 mg/kg rescued the *A. baumannii* infected larvae significantly (*p<0.001*). Polymyxin B at 10 mg/kg was used as positive control and PBS+PBS was used as trauma control (**Fig. 7B)**.

Finally, we tested the combination of MFQ+Polymyxin at 50 mg/kg of MFQ and 5 or 10 mg/kg of polymyxin B. Compared to standalone polymyxin B treatment at 5 mg/kg (**Fig. 7C**) or 10 mg/kg (**Fig. 7D**), the presence of MFQ at 50 mg/kg significantly improved survival in *A. baumannii*-infected larvae (*p < 0.001*)

## Discussion

MFQ was initially discovered in the late 1970s during research focused on finding effective treatments for malaria (22). Here, we show that MFQ is a bactericidal against *A. baumannii* even under acidic conditions (protonated), and drug-membrane interaction studies collectively showed that MFQ disrupts the bacterial cell membrane and decreases membrane fluidity. Membrane permeability assays in this study show that MFQ disrupts the bacterial membrane, enhancing permeability and antimicrobial activity. In this study, we explore the interaction of MFQ with the bacterial membrane. While other potential targets may exist, membrane disruption presents a clear and measurable effect in this context. Interestingly, MFQ exhibited synergistic effects with established antibiotics with activity as membrane permeabilizers, such as polymyxin B and colistin. The synergism of MFQ with polymyxin B observed across major Gram-negative pathogens such as *E. coli, B. pseudomallei, K. pneumoniae, P. aeruginosa*, and *E. aerogenes* can likely be attributed to their combined action on bacterial cell membranes. Polymyxin B disrupts the outer membrane of Gram-negative bacteria (23), and this could result in an increase in the membrane permeability and the entry of other antimicrobial agents, like MFQ, into the cell. Our working hypothesis is that the membrane permeabilizing effect may enhance the antimicrobial activity of MFQ, allowing it to exert an effect against these pathogens. Clinicians tend to use colistin more commonly than polymyxin B in many cases, though the choice depends on the specific clinical scenario, patient factors, and institutional preferences (24, 25). Overall, polymyxin B (26) is usually preferred for bloodstream infections (27) due to its more stable pharmacokinetics and was the primary agent used in this study. The MFQ+polymyxin B combination resulted in significant membrane damage in *A. baumannii*, highlighting the potential of MFQ when used in conjunction with other membrane permeabilizers.

The MFQ molecule consists of a quinoline ring system with a trifluoromethyl group attached to the aromatic ring and a piperidine ring-features that enhance the ability to traverse lipid membranes and interact with hydrophobic environments (28). We conducted structure-activity relationship studies on MFQ analogs lacking the piperidine moiety (through replacement with pyridinyl) or the piperidin-2-yl methanol (through replacement with piperidin-2-yl methanone) component and found a complete loss of antibacterial activity against *A. baumannii*, highlighting the critical role of the piperidin-2-yl methanol component in maintaining antibacterial activity. All-atom MD simulations at alkaline pH further demonstrated that piperidine moiety is the first component of MFQ to interact with the membrane. We also observed that the piperidine-containing moiety of MFQ interacts with the lipid bilayer to enhance upcoming MFQ uptake and improve membrane permeability contributing to its mode of its action.

Biofilms provide a protective environment for *A. baumannii*, significantly enhancing bacterial resistance to antimicrobial agents (29). The biofilm matrix acts as a physical barrier, limiting drug penetration (30). Additionally, bacteria within biofilms can adopt altered metabolic states, making them less susceptible to antibiotics (15). We found that MFQ inhibits *A. baumannii* biofilm formation and eradicates mature biofilms, consistent with previous studies (18, 31, 32) and investigated the impact of MFQ on biofilm-related gene expression. Next, we analyzed gene expression using RT-PCR with a 4-hour time point chosen for MFQ-treated *A. baumannii* to capture the early transcriptional response to treatment (33). Our gene-expression studies further explored that MFQ downregulated genes such as *csuA, csuB,* and *csuC* (encode components of a pili system involved in bacterial attachment to surfaces, which is a key initial step in biofilm formation), (34) as well as *pgaABCD*, which encodes poly-N-acetyl glucosamine (PNAG) (an extracellular matrix component that aids in biofilm stability and bacterial aggregation) (35). All of these down-regulated genes are critical for biofilm formation (30) further supporting the role of MFQ against biofilm.

Importantly, MFQ exhibited synergistic effects with the other membrane permeabilizer polymyxin B. We investigated the molecular interactions associated with the MFQ+polymyxin B combination using MD simulations at pH 7. We observed that polymyxin B (since it has 4 lysine residues that are protonated) first interacts electrostatically with the PG’s polar heads in the upper leaflet of the bilayer and moves through the membrane. The density distribution of the fatty acid tails, along with leucine (Leu) and phenylalanine (Phe) residues, suggests that hydrophobic residues primarily remain embedded at the interface between the lipid headgroups and tails, facilitating the insertion of polymyxin B into the bilayer. Our membrane interaction findings of the MFQ+polymyxin B combination indicate that polymyxin B increases the lipid area occupancy and decreases the hydrophobic thickness of the lipid tails, causing the lipid hydrophobic region to become more rigid. Being MFQ and polymyxin B protonated, it seems that the polymyxin B pushes MFQ to pass the membrane. This observation is key to understanding the mechanism by which the combination of MFQ+ polymyxin B works.

*A. G. mellonella* is a simple infection model (36) and also serves as a systemic toxicity testing model and found that MFQ improves the survival of infected larvae. Indeed, the MFQ+polymyxin B combination reduced the required dose of polymyxin B by 50 % compared to 10 mg/kg needed for standalone treatment in our experimental studies using *G. mellonella* model. These findings suggesting that MFQ might be more effective in decreasing the dose of toxic treatments such as colistin and polymyxin B that have been associated with severe side-effects such as hepatotoxicity, nephrotoxicity, and neurotoxicity (37). Further vertebrate model studies are needed to evaluate its pharmacokinetics, toxicity profile, and therapeutic efficacy. These studies will aid in determining the optimal dosing strategies to maximize antibacterial activity while minimizing potential side effects.

## Conclusion

MFQ is a membrane-active molecule effective against *A. baumannii* and major Gram-negative pathogens particularly when combined with polymyxin B. MFQ permeabilizes the bacterial membrane and reduces membrane fluidity. MD simulations and SAR studies reveal that the piperidine moiety is essential for the antibacterial activity of MFQ against *A. baumannii*. The strategy of improving MFQ-like analogs warrants further investigation. Novel MFQ analogs with structural similarities, particularly those incorporating the piperidine moiety, have the potential to enhance antibacterial activity against multidrug-resistant pathogens and improve *in vivo* efficacy, offering a promising strategy for combating infections caused by drug-resistant bacteria

## Materials and Methods

### Antibacterial susceptibility assays

Bacteria were all from the Mylonakis laboratory collection (**Table 1**). The broth microdilution method using Mueller-Hinton broth (MHB) was employed to assess the antibacterial activity of the test compounds as described previously. (38–40).

### Time to kill assays

The killing kinetics of MFQ, either alone or in combination at 4x MIC with other clinical agents, against *A. baumannii* (10^8 cells/mL) were analyzed (38, 40) using an exponential killing assay. Colony-forming units (CFUs) were enumerated after overnight incubation at 37°C. All assays were performed in duplicate.

### Membrane permeability and fluidity assays

Logarithmic *A. baumannii* cells with Sytox Green (at 0.0005 M for permeability assays) as described (38). Laurdan fluorescent dye (at 0.001 M for fluidity assays) was used as described previously (41). DMSO was included as the vehicle control.

### Biofilm assay

*A. baumannii* supplemented with glucose was used to monitor the biofilm inhibition and eradication of MFQ were followed as reported earlier(42).

### Real-Time PCR assay

*A. baumannii* was treated with sub-MIC of MFQ for 4h and RNA was isolated and gene expression analysis was performed as described in (39).

### Human blood cell (RBC) hemolysis assays

Human erythrocytes (4%) (Rockland Immunochemicals, Limerick, PA, USA) were used as described previously(38, 40).

### Checkerboard assays

Test compound A in 0.025 mL was arrayed horizontally and compound B was added vertically in a 8x8 combination followed by an addition of 0.05 mL of bacterial cells and incubated for 18h and growth inhibition was measured spectroscopically as described previously. (43).

### Computational Methods

Initially, we constructed a well-equilibrated (500 ns) membrane system to evaluate the free energy of drug permeation. Benzene was used as a control for this study. Both benzene and MFQ (∼7 Å) were randomly placed above two separate membrane systems. The system was then subjected to unbiased MD simulations for microseconds to assess permeability and the effect of the drug on the *A. baumannii* membrane.

#### System Preparation Details

To investigate the adsorption and permeation behavior of MFQ on the *A. baumanii* lipid bilayer model (44), MFQ was placed randomly at 7 Å distance from the top of the constructed membrane. Simulations were carried out along 6 different simulation systems (**Supplementary Table 1**). One system comprised only of the membrane and water, without any ligand, was utilized as the primary reference. The membrane consists of the following components: the main glycerophospholipids in the membrane are phosphatidylethanolamine (PE), phosphatidylglycerol (PG), acyl-phosphatidylglycerol (aPG) and cardiolipin (CL), along with monolysocardiolipin (MLCL) (45). The MFQ was orientated differently in the aqueous phase, with two distinct orientations at a 6.5 Å from the center of the PE/PG bilayer along the z-direction. For the MFQ (2MFQ/BL and 3MFQ/BL) molecules, they were positioned in the water phase within a range of 5 Å, with the hydrophobic aromatic ring remaining close to the polar head groups of the lipid bilayer. Similarly, the 4MFQ/BL inclusion complexes were placed on the polar head groups of the lipid membrane at 4.0 Å along the z-direction mentioned in **Supplementary Table 1**.

#### Simulation details

Two different approaches of molecular dynamics simulation were performed. The first study, which is run over six different systems, was conducted to investigate the behavior of the MFQ molecules or MFQ/polymyxin B in the lipid bilayer. The second simulation was set up for PMF calculations in systems containing bilayer along with a benzene (46) or MFQ molecule. The simulations were executed in the NPgT ensemble using AMBER 18 package (47). The AMBER14 lipid forcefield was used (20). The system was constructed using CHARMM-GUI builder (48). All simulations were carried out under temperature (323 K) and pressure (1 bar) to have the membrane above the melting point of individual lipids. The temperature was controlled by the Nose-Hoover thermostat (49) with a coupling time of 0.5 ps. The pressure control was done by coupling the simulation cell to a Parrinello-Rahman barostat (50), with coupling time constant 2 ps. All atom bond lengths were linked by using the LINCS algorithm (51). The equation of motion integrated by using the leap-frog algorithm with a time step of 2 fs. Electrostatic and van der Waals interactions cut-off were set at 12 Å. The long-range electrostatic interactions were treated using the particle mesh Ewald method (52). In all systems, unfavorable atomic contacts were removed by steepest descent energy minimization, prior to MD simulations. At first, while the positions of drugs were restrained, equilibration conducted in the NVT ensemble for 5 ns and then equilibration followed in NPT ensemble for 10 ns. After equilibration steps, all simulations were run for 800ns to 1 microsecond from their starting conditions and coordinates of the atoms.

#### PMF calculations

The free energy profiles including the potential of mean force (PMF) for translocation of MFQ (1-4 molecules) and benzene drug across the lipid bilayers were calculated using umbrella sampling followed by weighted histogram analysis method (WHAM) (53). Statistical errors were estimated using Bayesian bootstrap analysis (n=120) (54). To achieve initial configurations, the center of mass of a MFQ or benzene molecule were placed in the bilayer center, and then pulled through their center of mass into the bulk water along the z-axis using the umbrella sampling method (55). Drugs including MFQ and benzene were pulled along the bilayer (the z-axis) using a force constant of 1000 kJ mol^−1^ nm^−2^ and pulling rate of 0.01 nm ps^−1^ was used. A harmonic potential is exerted on the drug molecule by pulling, through which the center of molecule is transmitted at a specific rate. Among the snapshots of the simulated process of pulling along the z-axis, 31 windows (∼0.11 Å apart), ranging from the bilayer center (Z = ±0.0 nm) to the bulk water (Z = ± 3.5 nm), were selected. To avoid any drug-drug interactions, each window was explored in a separate simulation (10 ns run time each) and free energies were obtained from the last 2 ns, a smooth transition and good overlap between windows were obtained by using these conditions. Visualizations of systems have been created by using VMD (56) and PyMOL (57).

#### DFT calculations

Frames of the MFQ-BL complexes were extracted from the MD simulations snapshots. The observed interactions between drug-lipids were quantified at the DFT level in conjunction with the M06–2X (58) exchange–correlation functional and 6-311G (d,p) basis set (59) after full geometry optimization. The M06–2X functional was found to be a good descriptor for both covalent and non-covalent interactions (60). Solvent effects were estimated using Integral-Equation-Formalism Polarizable Continuum Model (IEF-PCM) (61). All calculations were carried out within the DFT algorithm by using the Gaussian16 B.01 package (62).

### Transmission electron microscopy

Mefloquine treated bacterial cells were fixed in 2.5% glutaldehyde in 0.1 M Na Cacodylate and 2% paraformaldehyde with 2mM calcium chloride 2-3 hours at 4°C and washed with cold 0.15 M Sodium Cacodylate buffer with 2 mM CaCl. Cells were treated with TCH, 20 min followed by a rinse for 5 times at 3 min interval using ddH2O at room temperature and treated with 2% OsO4 in ddH2O for 30 min at room temperature followed by an overnight treatment with in 1% uranyl acetate overnight at 4°C. Then cells were washed 5 times at 3 min interval using ddH2O at room temperature and cells were washed by Pb aspartate solution in 60° oven for 30 min and washed with 5 times at 3 min interval using ddH2O at room temperature. Cells were dehydrated in graded ethanol in 70%, 90%, 95%, 20 minutes each. Finally, cells were embedded in the resin followed by microscopy.

### In vivo assay

*Galleria Mellonella* wax moth model (n=16) was (36) used to evaluate the *in vivo* efficacy of MFQ or with polymyxin B combination as described previously {Peleg, 2009 #616}.

### Statistical analysis

All statistical analysis was carried out using GraphPad Prism version 10.02 (GraphPad Software, La Jolla CA, USA).

